# Transcriptional programs controlling lineages specification of mandibular epithelium during tooth initiation

**DOI:** 10.1101/2022.11.26.518052

**Authors:** Fan Shao, An-Vi Phan, Eric Van Otterloo, Huojun Cao

## Abstract

Tooth development starts with formation of the dental lamina, a localized thickening of the maxillary and mandibular epithelium, at specific location of oral cavity. How this dental lamina is specified within the continuous naïve maxillary and mandibular epithelium, remains as a critical and unresolved question. To identify potential genes and transcriptional regulatory networks that controlling dental lamina formation, we utilized single-cell Multiome-seq and Laser Microdissection coupled RNA-seq to profile gene expression and open chromatin of mandibular epithelium during tooth initiation. We comprehensively identified transcription factors (TFs) and signaling pathways that are differentially expressed among different domains (including dental lamina) along dorsal-ventral axis of mandibular epithelium. Specifically, we found *Sox2* and *Tfap2a/Tfap2b* forming two complementary domains along the dorsal-ventral axis of mandibular epithelium. Furthermore, dental lamina specific or enriched TFs such as *Pitx1, Pitx2* and *Zfp536* are presented around the interface of *Sox2* and *Tfap2a/Tfap2b*. We also identified correlation between signaling pathways and domain specific TFs. Through computational analysis of single-cell Multiome-seq dataset, we identified potential key/driver TFs and core transcriptional regulatory networks (TRNs) for lineages (oral, dental, skin) specification during tooth initiation. Our computational analysis also reveals a potential cross repression interactions between different groups of TFs (i.e. *Pitx1/Pitx2* and *Tfap2a/Tfap2b*) that might be essential for dental lamina specification.

## INTRODUCTION

The development of teeth is governed by reciprocal interactions between dental epithelium and the underlying neural crest derived mesenchyme. It has been shown that in most species, including mice and humans, tooth development starts with a localized thickening of the maxillary and mandibular epithelium, resulting in a ‘strip’ of dental epithelium (i.e., the dental lamina) that have the capability to induce a tooth development program(Jernvall and Thesleff, 2012). All teeth will originate within this dental lamina ‘strip’, going through a series of developmental stages including: placode, bud, cap, bell, etc. Classical tissue recombination experiments have shown that the tooth-inductive capacity, the odontogenic potential, first resides in the epithelium(Hu et al., 2014; Lumsden, 1988; Mina and Kollar, 1987). In mouse, the mandibular epithelium, before embryonic day 12.5 (E12.5), has the capability to induce tooth formation when recombined with secondary pharyngeal/branchial arch (2^nd^ PA) mesenchyme (odontogenic competent) but not with limb mesenchyme (odontogenic incompetent)(Zhang et al., 2005).

It is generally accepted that the segregation of maxillary and mandibular epithelium into a few distinct lineages (oral, dental and skin) is encoded by specific expression patterns of transcription factors (TFs) regulated by gradients of signaling pathways. Previous studies have shown that FGF and BMP pathways antagonize each other at the site of tooth initiation(Bei and Maas, 1998; Mitsiadis and Drouin, 2008; Neubüser et al., 1997; St.Amand et al., 2000; Stottmann et al., 2001). Additionally, a recent study shows SHH and BMP signal pathways are critical for establishment of oral-aboral (dorsal-ventral) axis of mandibular arch(Xu et al., 2019). Mouse knockout (KO) models of many genes expressed in the maxillary and mandibular epithelium before dental lamina formation—including *Pitx2, Sox2, Irx1, Lef1, Tbx1, Fgf8*, among others—have been generated and studied. However, despite tooth development defects at later stages, all models present with normal dental lamina formation(Andl et al., 2004; Balic, 2018; Bei, 2009; Cao et al., 2010; Dassule et al., 2000; De Moerlooze et al., 2000; Laurikkala et al., 2006; Lin et al., 1999; Liu et al., 2003; Mills et al., 1999; Mitsiadis and Drouin, 2008; Peters et al., 1998; Sasaki et al., 2005; Satokata and Maas, 1994; Trumpp et al., 1999; Yu et al., 2017; Yu et al., 2020). Thus, how this dental lamina is specified within the continuous naïve maxillary and mandibular epithelium, remains as a critical and unresolved question.

In this study, we applied Laser Microdissection (LMD) coupled with Smart-Seq2 protocol to generate transcriptome profiles of mandibular epithelium domains (including dental lamina) along dorsal-ventral axis. We comprehensively identified TFs and signaling pathways that are differentially expressed among different domains along dorsal-ventral axis of mandibular epithelium. Expression pattern of representative TFs were verified with immunofluorescent staining. We found *Sox2* and *Tfap2a/Tfap2b* forming two complementary domains along the dorsal-ventral axis of mandibular epithelium. Furthermore, dental lamina specific or enriched TFs such as *Pitx1* and *Pitx2* are presented around the interface of *Sox2* and *Tfap2a/Tfap2b*. Similarly, we found SHH pathway and WNT pathway forming complementary expression domains, with SHH pathway present at dorsal/posterior side of mandibular epithelium and WNT pathway present at ventral/anterior side of mandibular epithelium. In addition, we utilized 10x Genomics single cell Multiome-seq (sc-Multiome-seq) to simultaneously profile expression and open chromatin of the maxillary and mandibular epithelium at E10.5 (before dental lamina formation) and E12.5 (after dental lamina formation). Through computational analysis, we identified potential key/driver TFs and core transcriptional regulatory networks (TRNs) for lineages (oral, dental, skin) specification of maxillary and mandibular epithelium.

## RESULTS

### Transcriptome analysis of mandibular epithelium domains along dorsal-ventral axis

To identify genes that are critical for mandibular epithelium patterning and dental lamina formation, we first applied Laser Microdissection (LMD) coupled with Smart-Seq2 (Picelli et al., 2013) protocol (LMD-RNA-seq, **Fig. 1A**) to generate transcriptome profiles of four domains along the dorsal-ventral axis (also known as the oral-aboral axis) of the E11.5 mandibular epithelium. The four domains include: posterior epithelium (locate posterior of dental lamina), dental lamina, anterior epithelium (locate anterior of dental lamina) and aboral (ventral) epithelium (**Fig. 1B**). Bioanalyzer analysis show full length cDNA (peak around 2kb) and high quality sequencing library (peaks around 300-600bp) were generated by LMD-RNA-seq workflow.

**Figure. 1.**
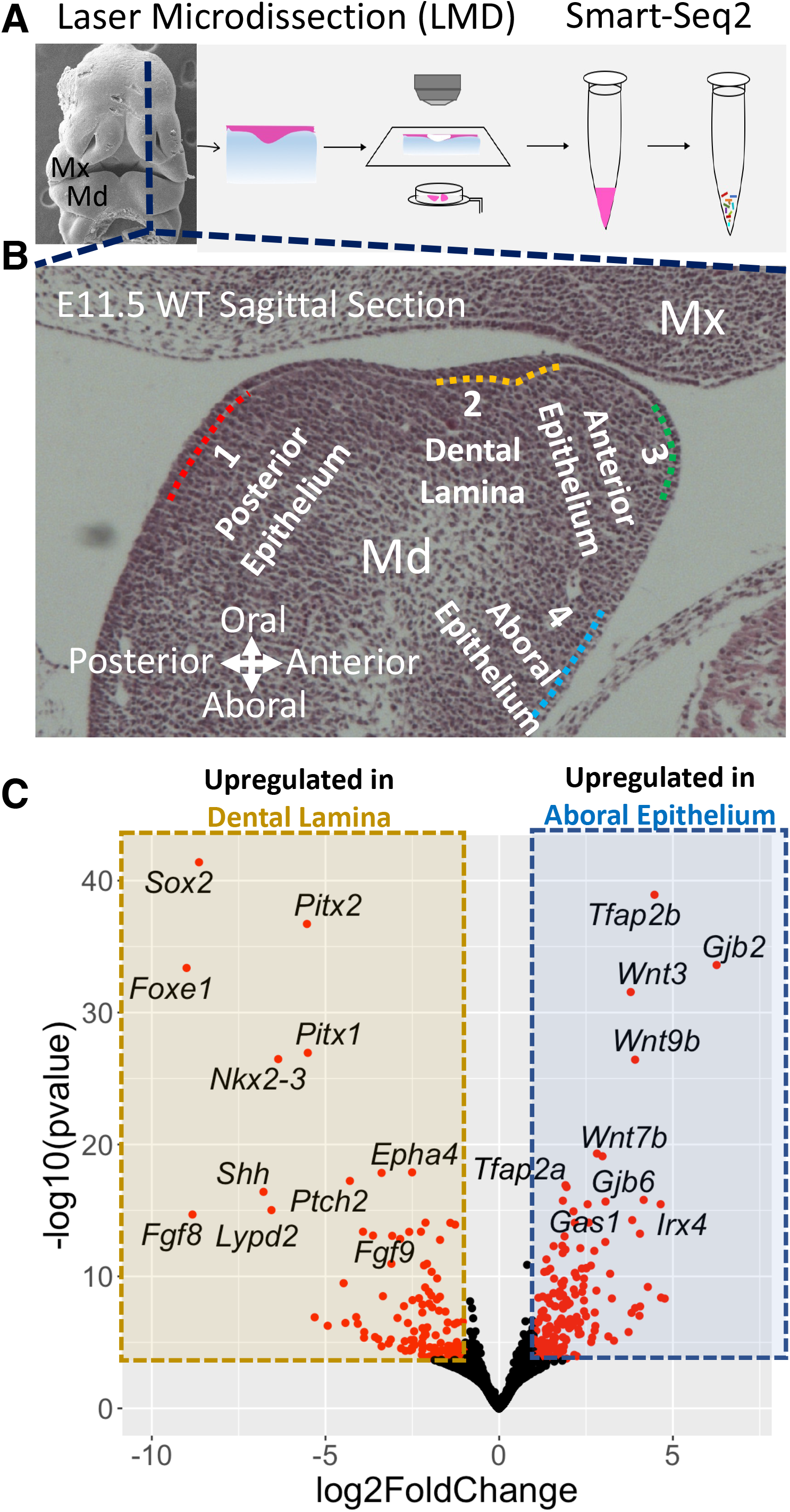
Transcriptome profiling with Laser microdissection (LMD) coupled with smart-seq2 protocol. **A**) Workflow of Laser microdissection (LMD) coupled with smart-seq2 protocol. **B**) LMD-RNA-seq libraries were prepared for 4 domains along dorsal - ventral axis of E11.5 mandible epithelium. **C**) Volcano plot for differentially expressed genes (DEGs) of E11.5 dental lamina comparing with aboral ectoderm. MxP: maxilla; MdP: Mandible

To identify domain specific or enriched genes, we performed pair-wise differentially expressed genes (DEGs) analysis of transcriptome profiles of these four domains with DESeq2(Love et al., 2014) (**Fig. 1C**). We found a large portion of top DEGs (based on P value and log2 fold change) are transcription factors (TFs) and signaling pathways genes. For example, in comparison of aboral epithelium with dental lamina, we found *Sox2, Pitx2, Foxe1, Pitx1, Nkx2-3, Shh, Fgf8*, etc. are among the most significantly upregulated (based on P value and log2 fold change) genes of dental lamina while *Tfap2b, Gjb2, Wnt3, Wnt9b, Wnt7b, Tfap2a, Irx4*, etc. are among the most significantly upregulated genes of aboral epithelium (**Fig. 1C**). To select genes that play major roles in mandibular epithelium patterning, we utilized a conserved filtration criteria for DEGs selection: adjusted P value should be less than 0.01 and absolute value of shrunken (ashr(Stephens, 2017) algorithm) log2 fold changes should be greater than 1.0. Based on this selection criteria, we comprehensively identified domain specific genes for the four domains along the dorsal-ventral axis of the E11.5 mandibular epithelium.

### Expression pattern of domain specific transcription factors in mandibular epithelium along dorsal-ventral axis

To visualize the expression pattern of domain specific TFs, we generated a expression heatmap with abovementioned LMD-RNA-seq transcriptome data (**Fig. 2**). From the heatmap of domain specific TFs and transcriptome profiles, we found anterior epithelium and aboral epithelium are molecularly similar to each other, expressing the largely same set of TFs: *Tfap2a, Tfap2b, Msx1, Msx2, Dlx3, Lef1*, etc. Similarly, posterior epithelium and dental lamina share some TFs including: *Sox2, Pitx2, Nkx2-3, Pitx1, Isl1, Foxe1, Tbx1*, etc. However, we found one group of TFs are more specifically restricted to dental lamina: *Zfp536, Ascl4, Sp6, Ascl5, Irx1*, etc, while another group of TFs are specific for posterior epithelium: *Foxa1, Foxa2, Sox5*, etc. Some of the dental lamina specific TFs (e.g. *Pitx2, Sox2, Tbx1*, etc.) are known to have critical roles during tooth development(Arnold et al., 2011; Cao *et al*., 2010; Caton et al., 2009; Juuri et al., 2013; Juuri et al., 2012; Lin *et al*., 1999; Liu *et al*., 2003; Lu et al., 1999; Sun et al., 2016; Yu *et al*., 2020), while for others TFs (e.g. *Foxe1, Nkx2-3, Klf5, Zfp536*, etc.), their function, specifically during mandibular epithelium patterning and dental lamina formation, is unclear. Specifically, *Pitx2* is one of the earliest and most specific markers for dental epithelium, and if removed, causes tooth development arrest at placode (maxillary teeth) or bud (mandibular teeth) stage with full penetrance(Balic, 2018; Lin *et al*., 1999; Liu *et al*., 2003; Lu *et al*., 1999; Mucchielli et al., 1997; Thesleff, 2008). Previous studies have shown that *Pitx2* expression is presented at broader domain in oral cavity before dental lamina formation, and as development proceeded, the expression of *Pitx2* become more confined to the dental epithelium (St.Amand *et al*., 2000). Importantly, lineage tracing experiments have shown that although *Pitx2* expression become progressively restricted to dental epithelium, *Pitx2* daughter cells contribute to both oral epithelium and facial skin epithelium (these cells were originated from *Pitx2* positive cells and gradually lost expression of *Pitx2*)(Liu *et al*., 2003).

**Figure. 2.**
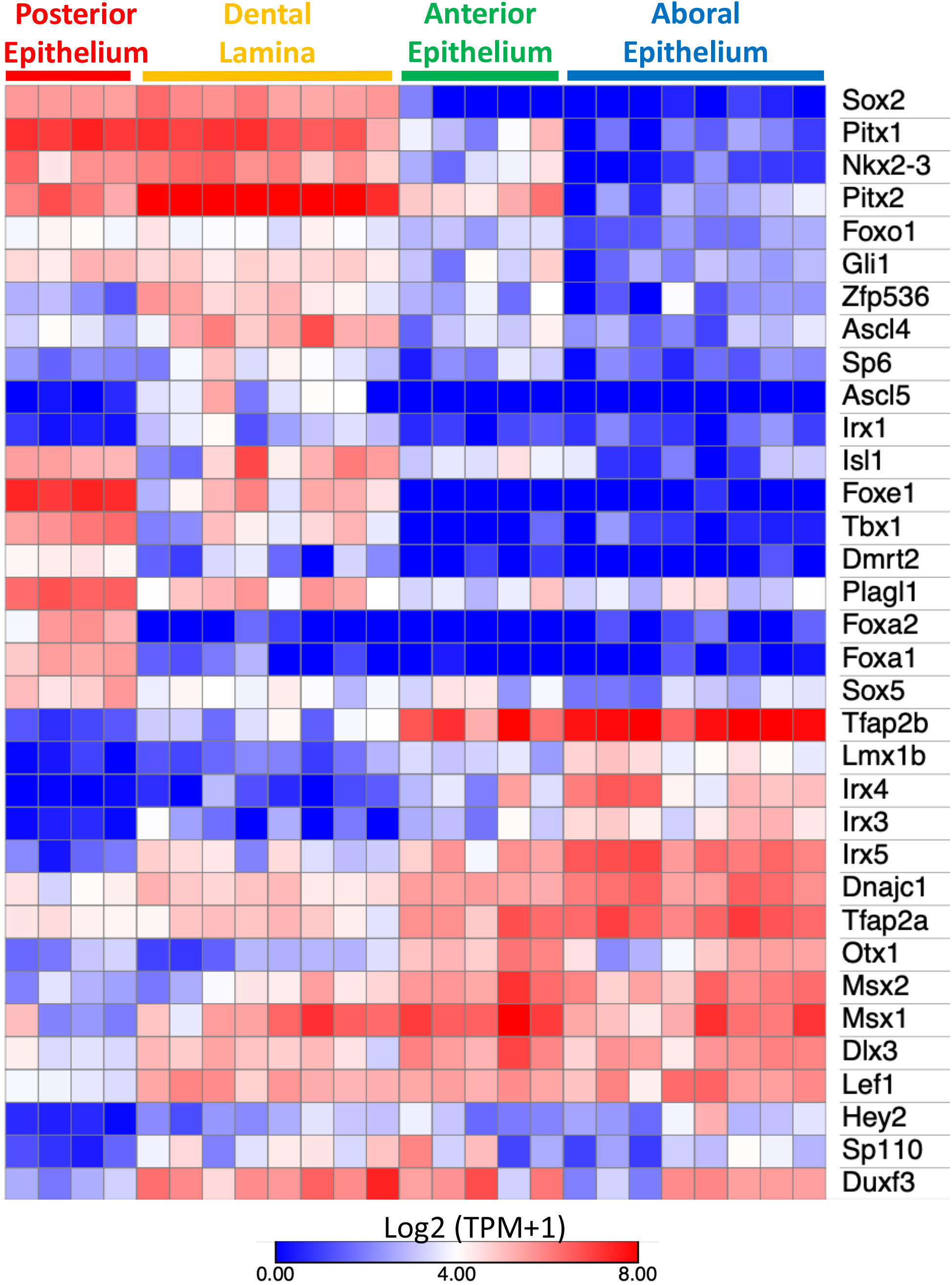
Expression heatmap of domain specific transcription factors. Colors are based on log2 transformed (TPM +1) values. TPM: transcript per million

To verify expression pattern of these domain specific TFs, we utilized immunofluorescent staining (IF) to directly visualize expression pattern of representative TFs (including: SOX2, PITX2, TFAP2A, TFAP2B and LEF1) along dorsal-ventral axis of mandibular epithelium at E9.5 and E11.5 (**Fig. 3**). Overall, we found that IF staining (protein level) results generally agree with LMD-RNA-seq result (mRNA level). Broadly, we found SOX2 and TFAP2A/TFAP2B forming two complementary domains along the dorsal-ventral axis of mandibular epithelium: with SOX2 locate at dorsal side while TFAP2A/TFAP2B locate at ventral side. PITX2 and LEF1 expression are presented in the middle, around the interface of SOX2 and TFAP2A/TFAP2B (**Fig. 3**). Specifically, at E11.5, PITX2 expression cover the whole dental lamina (**Fig. 3G)** and LEF1 expression is restricted to anterior part of dental lamina (**Fig. 3F**). At E11.5, SOX2 expression is presented at pharyngeal endoderm, oral epithelium and posterior part of dental lamina (**Fig. 3A & 3D)**. SOX2 expression is quickly reduced in the anterior part of dental lamina and didn’t extend beyond the dental lamina (**Fig. 3C & 3F**). On the other hand, at E11.5, TFAP2A/TFAP2B expression is presented at ventral side of mandibular epithelium and quickly reduced after reaching SOX2 positive cells in dental lamina (**Fig. 3B & 3H)**. Interestingly, the expression pattern of SOX2, TFAP2A, TFAP2B and LEF1 seems broader at E9.5 (before dental lamina formation) than at E11.5, having broader overlapping domains, suggesting a progressive restriction or enrichment of expression of these domain specific TFs.

**Figure. 3.**
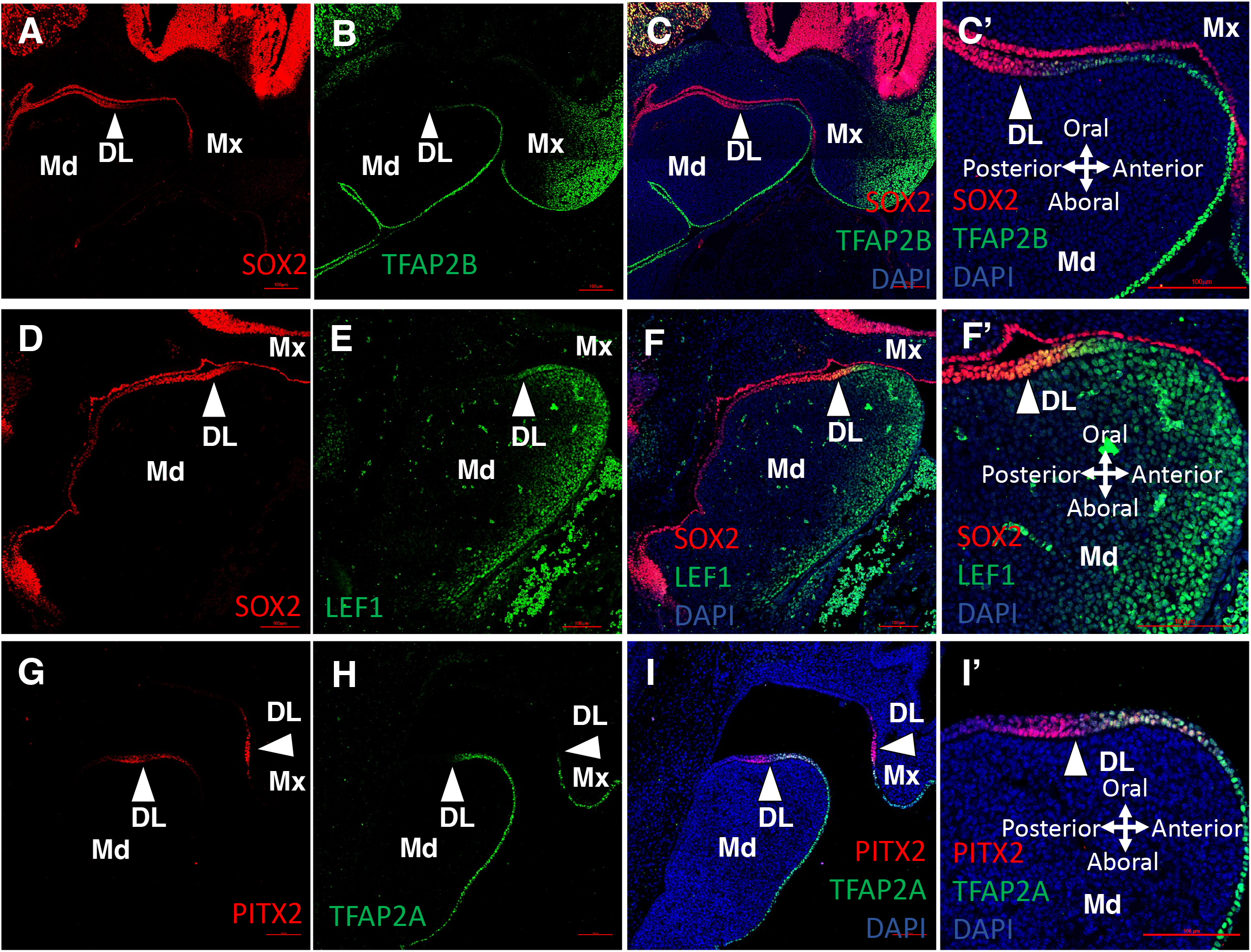
Expression pattern of selected domain specific transcription factors. **A**,**B**,**C**) SOX2, TFAP2B and merged immunofluorescence (IF) staining in E11.5 mouse heads. **D**,**E**,**F**) SOX2, LEF1 and merged IF staining in E11.5 mouse heads. **G**,**H**,**I**) PITX2, TFAP2A and merged IF staining in E11.5 mouse heads. **C’, F’, I’**) Enlarged images of dental lamina regions in **C, F, I**. Mx: Maxilla; Md: Mandible; DL: dental lamina; DAPI: DAPI DNA staining.

### Single-cell RNA-seq & ATAC-seq analysis of the E10.5 and E12.5 mandibular and maxillary epithelium

To further delineate the process of mandibular epithelium patterning and dental lamina formation, we utilized 10x Genomics single cell Multiome-seq (sc-Multiome-seq) to simultaneously profile expression and open chromatin of the first pharyngeal arch (1^st^ PA: mandibular & maxillary processes) epithelium at E10.5 (5K cells) and E12.5 (6K cells) at single cell resolution. We separated 1^st^ PA epithelium from the underlying mesenchyme with Dispase II protocol (Otterloo et al., 2022). At E12.5, we found at tooth placode sites, epithelium and mesenchyme formed strong connection that cannot be completely disrupted with our brief Dispase II treatment (20 minutes at 37°C). Accordingly, in our initial analysis of sc-Multiome-seq data, we found a small portion of cells in E12.5 dataset that are mesenchymal cells. These mesenchymal cells express *Pax9, Bmp4* and *Lef1*, suggesting they are dental mesenchymal cells. These mesenchymal cells were removed from current analysis.

Unsupervised Weighted-Nearest Neighbor (WNN) clustering with Seurat package(Hao et al., 2021) of the remaining epithelial cells revealed 10 cell clusters and 16 cell clusters in E10.5 and E12.5 1^st^ PA epithelium datasets, respectively (**Fig. 4**). We identified markers for each cluster and visualize the expression of top clusters markers with heatmap. Many of these markers are domain specific genes we identified in the abovementioned LMD-RNA-seq analysis (**Fig. 2**). Based on the expression of known markers, we mapped clusters of E10.5 and E12.5 1^st^ PA epithelium datasets to domains of mandibular and maxillary epithelium along dorsal-ventral axis. Specifically, in E10.5 dataset, clusters 8 and 3 represent dorsal/posterior epithelium domain, expressing *Sox2, Foxe1, Foxa1, Foxa2* and *Shh*; clusters 7, 5, 1, 2 and 10 represent ventral/anterior domain, expressing *Tfap2a, Tfap2b, Msx1, Irx4* and *Cxcl14;* clusters 6, 9, 4 and 0 represent the middle domain, expressing *Pitx1, Pitx2, Zfp536* and *Irx1* (**Fig. 4A & 4B**). Similarly, 16 clusters in E12.5 dataset can be mapped to these 3 major domains (dorsal/posterior, middle and ventral/anterior) based on expression of similar set of domain specific markers (**Fig. 4C & 4D**). Interestingly, we noticed that the spatial organization of clusters of E10.5 and E12.5 1^st^ PA epithelium in the 2D UMAP plot seems correlated well with their in vivo physical spatial organization along dorsal-ventral axis. Specifically, in 2D UMAP plot, we found *Sox2* and *Tfap2a/Tfap2b* formed two complementary domains, with dental epithelium markers including *Pitx1, Pitx2, Zfp536* and *Irx1* expressed in middle clusters (**Fig. 4**).

**Figure. 4.**
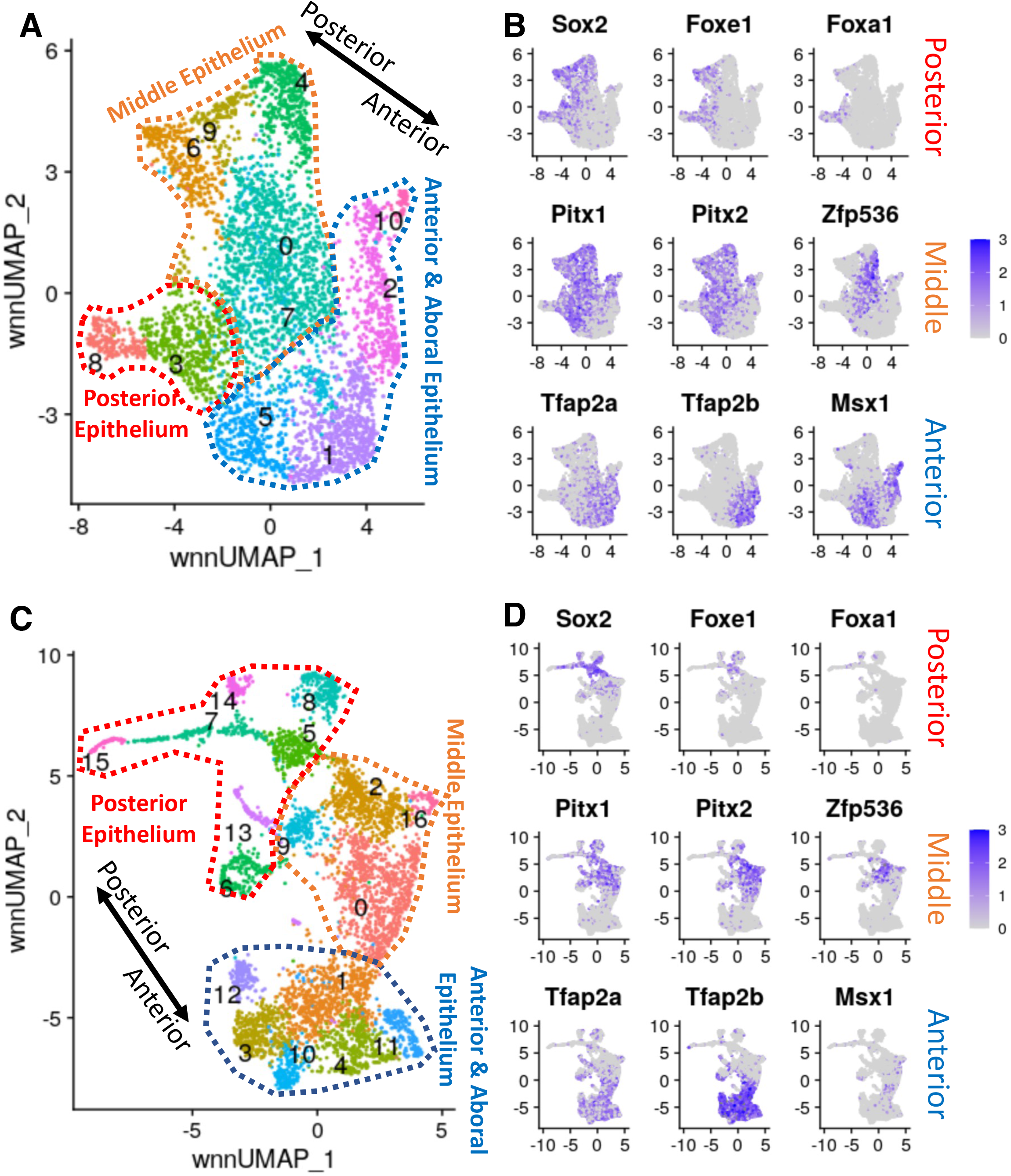
Single cell Multiome-seq of E10.5 and E12.5 maxillary and mandibular epithelium. **A**,**C)** UMAP plot showing three major groups of cells of E10.5 and E12.5 maxillary and mandibular epithelium. **B**,**D)** Expression Feature plots of representative domain specific transcription factors of E10.5 and E12.5 maxillary and mandibular epithelium.

Based on LMD-RNA-seq transcriptome profiles, IF staining and sc-Multiome-seq results, during dental lamina formation, we can broadly partitioned mandibular epithelium into 3 major domains along dorsalventral axis: dorsal/posterior domain expressing *Sox2, Foxe1, Foxa1, Foxa2*, etc.; ventral/anterior domain expressing *Tfap2a, Tfap2b, Msx1, Irx4*, etc.; and finally, the middle domain expressing *Pitx1, Pitx2, Zfp536, Irx1, etc*. It is also worth mentioned that some of these domain specific TFs formed largely overlapping pattern with slightly difference. For example, both *Pitx1* and *Pitx2* are mainly expressed in middle domain marking dental epithelium, however, expression of *Pitx1* extend more posteriorly than *Pitx2* (**Fig. 2, Fig. 4**). Similarly, the expression of *Tfap2a* and *Tfap2b* largely overlapped within ventral/anterior domain, however, the expression of *Tfap2a* is broader than *Tfap2b* beyond ventral/anterior domain (**Fig. 2** and **Fig. 3**).

### Correlation between domain specific transcription factors and signaling pathways

In both LMD-RNA-seq transcriptome profiles and sc-Multiome-seq results, we found expression of many signaling pathways genes are correlated with domain specific TFs. For example, in comparison of aboral/ventral epithelium with dental lamina, we found expression of *Fgf8, Fgf9, Shh, Ptch2* and *Epha4* in dental lamina and expression of *Wnt3, Wnt9b, Wnt7b* and *Gas1* in aboral/ventral epithelium. Similar as TFs, we comprehensively identified domain specific signaling pathways genes (based on KEGG pathway database(Kanehisa and Goto, 2000)) with abovementioned LMD-RNA-seq dataset. We generated expression heatmap for all genes that are significaltly differentially expressed in abovementioned pair-wise DEG analysis (**Fig. 1**) and are included in KEGG pathway database.

We found expression of dorsal/posterior domain TFs (*Sox2, Foxe1, Foxa1, Foxa2*, etc.) correlate with several genes of SHH pathway. Specifically, we found expression of *Shh* is restricted to dorsal side of E11.5 mandibular epithelium including dental lamina, it is absent in ventral side of mandibular epithelium. This result agrees with previous studies(Cobourne et al., 2004; Sarkar et al., 2000; Xu *et al*., 2019). Additionally, we found expression of receptors (*Ptch1* and *Ptch2*) and downstream transcription factor (*Gli1*) of SHH pathway are also enriched in dorsal side of mandibular epithelium. Interestingly, we found expression of *Gas1*, a negative regulator of SHH pathway, is restricted to ventral side of mandibular epithelium (**Fig. 5A**). Similarly, in E10.5 1^st^ PA epithelium sc-Multiome-seq dataset, we found restriction of SHH ligand (*Shh*), receptors (*Ptch1* and *Ptch2*), and downstream transcription factors (*Gli1* and *Gli3*) in posterior and middle domains, while expression of *Gas1* is enriched in anterior domain (**Fig. 5B**).

**Figure. 5.**
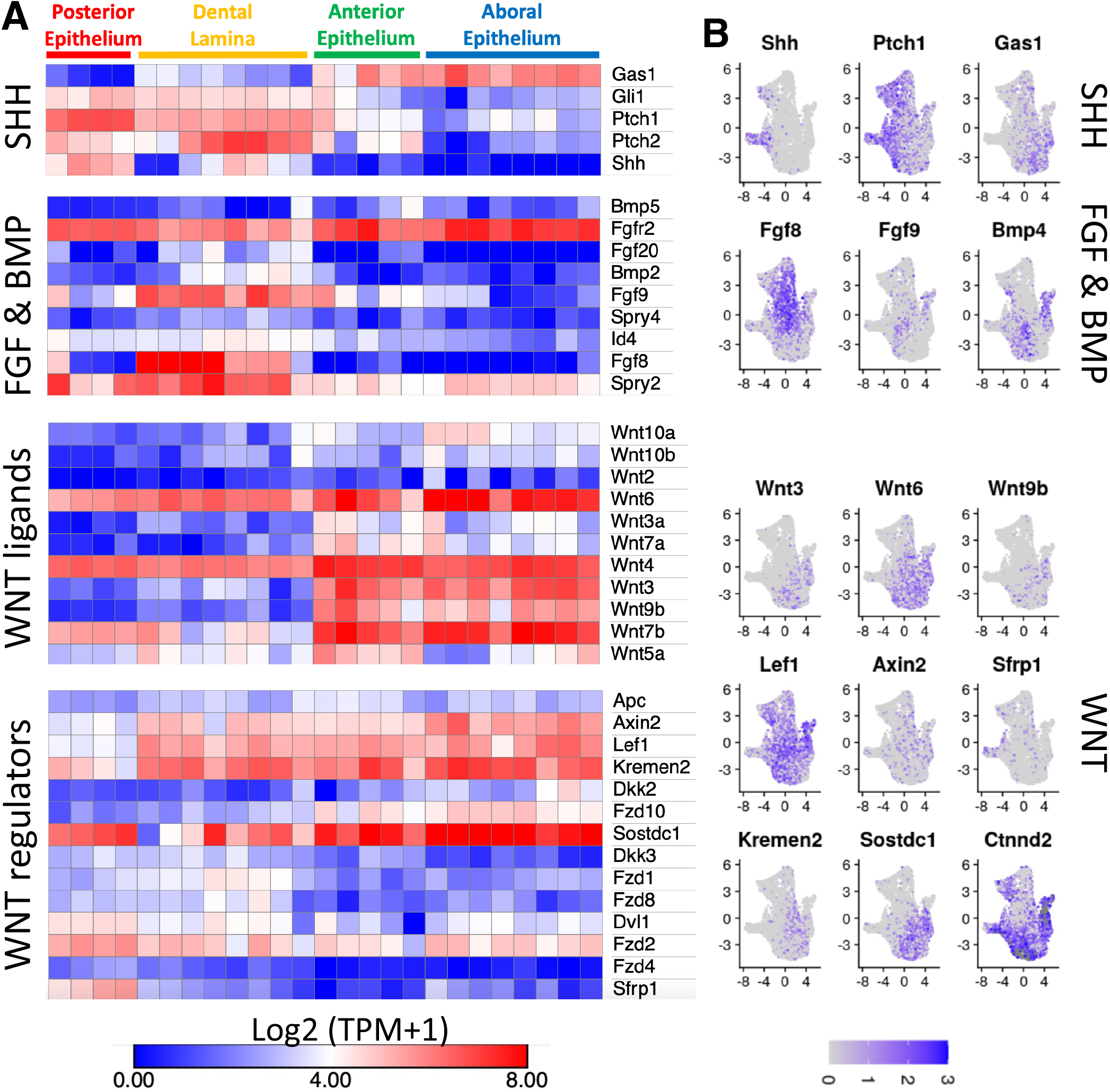
Expression pattern of differentially expressed signaling pathway genes. **A**) Expression heatmap of differentially expressed signaling pathway genes. Colors are based on log2 transformed (TPM +1) values. **B**) Expression feature plots of selected differentially expressed signaling pathway genes of E10.5 maxillary and mandibular epithelium. TPM: transcript per million

We found expression of ventral/anterior domain TFs (*Tfap2a, Tfap2b, Msx1, Irx4*, etc.) correlate with multiple genes of WNT pathway. We found expression of multiple WNT ligands (*Wnt3a, Wnt3, Wnt7a, Wnt9b, Wnt10a* and *Wnt10b*) are restricted to ventral/aboral side of E11.5 mandibular epithelium (**Fig. 5A**). Few other WNT ligands (*Wnt6, Wnt4, Wnt7b* and *Wnt5a*) are expressed in dental lamina and posterior epithelium, but their expression is significantly upregulated in ventral/aboral epithelium. Interestingly, we found several negative regulators of WNT (*Axin2, Kremen2*, and *Sostdc1)* are highly expressed in ventral/aboral epithelium. Accordingly, closely examination of LEF1 staining, a readout of WNT pathway activity, revealed that the highest level of LEF1 is located at anterior part of dental lamina, it’s level become gradullay diminished moving towards ventral/aboral end of epithelium (**Fig. 3E, 3F**). However, the underlying mesenchyme of ventral/aboral side of epithelium express high level of LEF1, suggesting that WNT ligands secerted by ventral/aboral side of epithelium induce WNT pathway activity in the underlying mesenchyme (**Fig. 3E, 3F)**. Previous studies have shown that constitutive activation of WNT pathway in oral and dental epithelium leads to supernumary tooth formation(Järvinen et al., 2006; Liu et al., 2008; Wang et al., 2009; Zhou et al., 2019). The expression pattern of these negative regulators of WNT pathways might be critical in repress dental epithelium fate in ventral/aboral epithelium. In E10.5 1^st^ PA epithelium sc-Multiome-seq dataset, we found another potential negative regulator of WNT pathway, *Ctnnd2(Turner et al., 2015)*. *Ctnnd2* is highly expressed in clusters of both dorsal/posterior and ventral/aboral domains but are dramatically reduced in clusters of middle domain.

In summary, similar as domain specific TFs, we found SHH pathway and WNT pathway forming complementary expression domains, with SHH pathway present at dorsal/posterior side of mandibular epithelium and WNT pathway present at ventral/anterior side of mandibular epithelium. Specifically, at E11.5, we found activity of SHH pathway (*Ptch1, Ptch2* and *Gli1*) and WNT pathway (*Lef1* and *Axin2*) are converged in dental lamina (**Fig. 5A**). Additionally, we found expression of multiple genes in FGF and BMP pathways are specifically restricted or enriched in dental lamina, including *Fgf20, Bmp2, Fgf9, Fgf8, Id4*, etc (**Fig. 5A**).

### Prediction of key transcription factors and transcriptional regulatory networks for dental epithelium

We integrated scRNA-seq and scATAC-seq to identify potential key TFs for cell clusters in E10.5 and E12.5 1^st^ PA epithelium sc-Multiome-seq dataset. Specifically, we utilized chromVAR package to identify TFs’ motifs associated with variability in chromatin accessibility between individual cells(Schep et al., 2017). We then use PRESTO package to perform fast differential analysis with gene expression values and motif variability values calculated with chromVAR(Hao *et al*., 2021; Schep *et al*., 2017). We ranked all TFs based on “AUC” statistic calculated by PRESTO package. Top 10 predicted key TFs for clusters of E10.5 1^st^ PA epithelium were shown in **Fig. 6A**. We found *Foxa1, Foxe1* and multiple *Sox* family TFs rank at top for clusters of oral epithelium lineage (dorsal/posterior domain); *Pitx1, Pitx2, Sox2*, WNT pathway downstream TFs and SHH pathway downstream TFs rank at top for clusters of dental epithelium lineage (middle domain); and *Tfap2a, Tfap2b, Msx1, Msx2* and *Dlx3* rank at top for clusters of skin epithelium lineage (ventral/anterior domain).

**Figure. 6.**
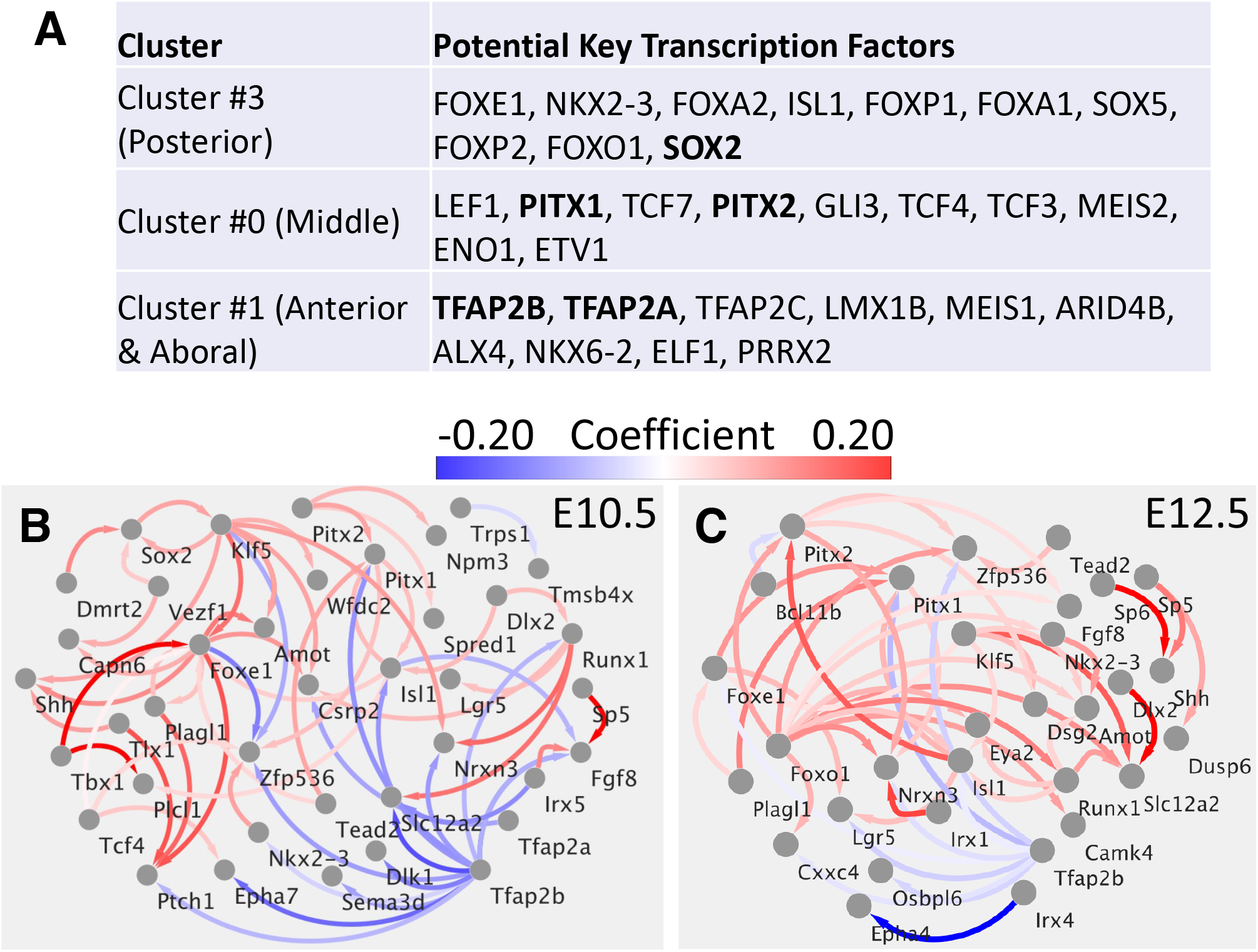
Inferred core Transcriptional Regulatory Networks for dental epithelium. **A**) Computationally inferred top driver transcription factors. **B)** Inferred core Transcriptional Regulatory Networks for dental epithelium at E10.5 and **C)** at E12.5. The arrow linked TFs to its targets. The arrow was colored based on its fitted coefficient in linear regression model.

We leveraged our sc-Multiome-seq data (simultaneous profiling of gene expression and chromatin accessibility in individual cells) to computational infer transcriptional regulatory networks (TRNs) regulating the cell lineages specification during dental lamina formation. Briefly, **I)** we first identified markers of dental lineage clusters (middle domain) and skin lineage clusters (ventral/anterior domain); **II)** we utilized Signac(Stuart et al., 2021) package to link potential cis-regulatory elements (CREs) to dental/skin domain specific genes; **III)** we scanned CREs for transcription factor motifs; **IV)** we utilized Pando(Fleck et al., 2021) package to build a linear regression model for the expression of dental/skin genes based on associated CREs and transcription factor activity. The fitted coefficient of transcription factor can be interpreted as quantitative measurement of transcription factor effect on the target genes, with positive coefficient means activation and negative coefficient means repression. Only significant interactions (adjusted p value < 0.01) were kept in the TRNs. **V)** finally, we leveraged our LMD-RNA-seq results to narrow down core TRNs for dental and skin epithelium development. The inferred core TRNs revealed that *Pitx1, Pitx2, Sox2, Klf5, Foxe1* etc. driving dental epithelium development while *Tfap2a, Tfap2b, Irx4* and *Irx5* negatively regulating the core dental TRNs (**Fig. 6B**). Conversely, we found *Tfap2a, Tfap2b, Irx4* and *Irx5* driving core skin TRNs while *Pitx1* and *Pitx2* repress it. Additionally, the inferred TRNs suggest the within-group TFs activation and cross-group TFs repression might be important for the dental/skin lineage specification. For example, at E10.5 *Sox2* and *Klf5* active each other; *Tfap2a* and *Tfap2b* active each other at both E10.5 and E12.5; *Tfap2b* repress *Pitx1* expression at both E10.5 and E12.5 (**Fig. 6B**).

## DISCUSSION

In this study, we utilized two transcriptome profile technologies: Laser Microdissection (LMD) coupled RNA-seq and single cell Multiome-seq (scRNA-seq & scATAC-seq). Both technologies have its advantages and limitations. Specifically, LMD-RNA-seq enable us to profile cells at specific locations. scRNA-seq, on the other hand, give us a higher resolution view. However, the detection sensitivity of scRNA-seq is lower than LMD-RNA-seq. For genes with low or moderate expression level, LMD-RNA-seq provide more accurate results than scRNA-seq. For example, LMD-RNA-seq enable us to detect expression pattern of *Wnt10a* and *Wnt10b*. In examination of scRNA-seq, we found expression values of *Wnt10a* and *Wnt10b* in most cells become zero.

Several recent scRNA-seq profile of the mandibular arch have been published: 1) scRNA-seq of the E10.5 mandibular arch; 2) scRNA-seq of E10.5, E12.5 and E14.5 maxillary and mandibular arch; 3) scRNA-seq of E12.5 mandibular epithelium (Xu *et al*., 2019; Ye et al., 2022; Yuan et al., 2020). Our sc-Multiome-seq profile of E10.5 and E12.5 maxillary and mandibular epithelium enable us to simultaneously measure expression and open chromatin, to better understanding of transcriptional regulatory networks. It is also particular interesting to integrate multiple datasets together to infer communication networks between epithelium and mesenchyme (epithelium account less than 10% of mandibular arch).

In this study, we found it is very common that paralogs (*Foxa1* and *Foxa2*; *Pitx1* and *Pitx2*; *Tfap2a* and *Tfap2b*; etc.) share similar expression patterns. It is very likely that genetic redundancy provided by paralogs contribute to the lack of models with disrupted dental lamina formation. It will be interesting to generate and investigate double or triple knockout mouse models. For example, single knocking-out *Tfap2a* or *Tfap2b* with an early ectodermal CRE line (CRECT(Schock et al., 2017), active ∼ E7.5) did not result in any obvious defects in tooth development. However, knockout of both *Tfap2a* and *Tfap2b* together leads to ectopic incisor that formed at the aboral (ventral) side of the mandible(Woodruff et al., 2021).

There are several limitations of current study. Our computational analysis predicts that *Tfap2a* and *Tfap2b* repress dental lineage networks while *Pitx1* and *Pitx2* inhibit skin lineage networks. The cross repression between different group of TFs might be essential to convert broad pattern domains setup by gradients of signaling pathways to sharp and precise domains. However, currently, this cross repression is solely based on correlation of gene expression and open chromatin with wild type mouse cells. Further studies with knockout mouse models will be necessary to test these computational predictions. Ideally, direct measurement of DNA binding by these domain specific TFs (such as *Pitx1, Pitx2, Tfap2a* and *Tfap2b*) with CUT&RUN assay should be performed to identify direct targets of these TFs during dental lamina formation. Similarly, the correlation between signal pathways and TFs observed in this study need perturbation experiments for validation.

## MATERIALS AND METHODS

### Histology, H&E and immunofluorescence

Mouse embryos were harvested and washed with ice-cold 1× PBS and fixed with 4% paraformaldehyde in PBS for 4 hours. Following fixation, samples were dehydrated through a graded ethanol series, embedded in paraffin wax and sectioned (7 μm). Standard Hematoxylin and Eosin (H&E) staining was used to examine tissue morphology. For immunohistochemistry and immunofluorescence, sections were incubated in citrate buffer (pH=6.0) in a 100°C water bath for 20 min or autoclaving in 0.1 M Tris-HCl buffer (pH 9.0) for 5 minutes. Sections were then blocked with 10% goat serum and incubated with primary antibodies overnight in 4°C. Following primary antibodies (αTFAP2A, Abcam ab108311; αTFAP2B, Abcam ab221094; αLEF1, Cell Signaling, 2230; αSOX2, R&D Systems, AF2018; αPITX1, Abcam ab70273; and αPITX2, R&D Systems, AF7388) were diluted in TBS (0.5% Tween in phosphate-buffered saline) containing 0.1% Triton X-100, 5% goat serum and 1% BSA. Then sections were washed with PBS, incubated with secondary antibodies, and stained with DAPI. Images were captured using a Nikon Eclipse microscope or a Zeiss 700 confocal microscope.

### Laser Microdissection (LMD) coupled RNA-seq

Laser Microdissection were performed with Leica LMD7000 Laser Microdissection Systems. For preparation of sections, Methacarn fixation solution was used instead of 4% paraformaldehyde. Cells were cut at 20 × magnification while keeping laser power to a minimum. After cells were collected in cap, tubes were spin down quickly and 5 μl of Smart-seq2 lysis buffer (with 0.5% Triton X-100, 5μg Protease K and 2 U μl^−1^ RNase inhibitor) was added, followed by pipetting up and down to mix. Cells were lysed for 1 hour at 60°C. Then samples were treated with 1U DNase I for 30 min. Standard Smart-seq2 protocol were followed for cDNA preparation, Tn5 Tagmentation and next generation sequencing libraries preparation. Quality of cDNA and sequencing libraries were analyzed with bioanalyzer and real time PCR.

### RNA-seq data analysis

RNA-seq reads were quality checked using the FastQC tool (http://www.bioinformatics.babraham.ac.uk/projects/fastqc). Low-quality and adapter sequences were removed with the Trimmomatic(Bolger et al., 2014). Expression of transcripts was quantified using the Salmon tool(Patro et al., 2017), and estimates of transcript abundance for gene-level analysis were imported and summarized using the tximport(Soneson et al., 2015) function of the R/Bioconductor software suite(Huber et al., 2015). Differentially expressed genes (DEGs) were identified by applying the R/Bioconductor package DeSeq2(Love *et al*., 2014). Heatmap were generated with Morpheus (https://software.broadinstitute.org/morpheus).

### Single-cell Multiome-seq and data analysis

E10.5 and E12.5 first pharyngeal arch epithelium were separated from mesenchyme with brief Dispase II (2mg/ml in PBS) treatment (20 mins @ 37°C) as previous described (Otterloo *et al*., 2022). Single-Cell were prepared with cold protease protocol as previous described(Denisenko et al., 2020; Sekiguchi and Hauser, 2019). Single nucleus were prepared with Chromium Nuclei Isolation Kit. 0.4% trypan blue was used for quality check. 10x Genomics single cell Multiome-seq (sc-Multiome-seq) platform were used to profile expression and open chromatin.

Cellranger-arc was used to process sequencing reads. Specifically, cellranger-arc count was used to align sequencing reads with mm10 reference genome and generate feature counts matrix for expression and open chromatin. We then utilized Seurat package(Hao *et al*., 2021) to import the feature counts matrix of expression and open chromatin. Standard Seurat workflow was used for filtration of low-quality cells, data normalization, dimension reduction. We also use seurat package to regress out effects of cell cycle and sex. We utilized MACS2 package(Zhang et al., 2008) to identify peaks of scATAC-seq data. Weighted Nearest Neighbor analysis was performed for unsupervised clustering. Signac package(Stuart *et al*., 2021) was used to compute the correlation between gene expression and accessibility of nearby peaks. chromVAR package(Schep *et al*., 2017) was used to compute motif accessibility score for individual cell.

## ACKNOWLEDGEMENTS

We are grateful to the Central Microscopy Research Facility (University of Iowa) for providing us training and support of Laser Microdissection experiments. We are grateful to the Iowa Institute of Human Genetics (IIHG) Genomics Division (University of Iowa) for helping us with single-cell Multiome-seq. We also thank University of Iowa Research Services for providing High Performance Computing Resources. Additionally, we would like thank members of the Cao, Amendt and Van Otterloo laboratories for helpful discussion and suggestions. This study was supported by grants from National Institute of Dental and Craniofacial Research (R21DE029828, R03DE028354) and seed grant from University of Iowa College of Dentistry and Dental Clinics.

